# Cannabis Use in Youth is Associated with Limited Alterations in Brain Structure

**DOI:** 10.1101/443911

**Authors:** J. Cobb Scott, Adon F. G. Rosen, Tyler M. Moore, David R. Roalf, Theodore D. Satterthwaite, Monica E. Calkins, Kosha Ruparel, Raquel E. Gur, Ruben C. Gur

## Abstract

Frequent cannabis use during adolescence has been associated with alterations in brain structure. However, studies have featured relatively inconsistent results, predominantly from small samples, and few studies have examined less frequent users to shed light on potential brain structure differences across levels of cannabis use. In this study, high-resolution T1-weighted MRIs were obtained from 781 youth aged 14-21 years who were studied as part of the Philadelphia Neurodevelopmental Cohort. This sample included 147 cannabis users (109 Occasional [≤1-2 times per week] and 38 Frequent [≥ 3 times per week] Users) and 634 cannabis Non-Users. Several structural neuroimaging measures were examined in whole brain analyses, including gray and white matter volumes, cortical thickness, and gray matter density. Established procedures for stringent quality control were conducted, and two automated neuroimaging software processing packages were used to ensure robustness of results. There were no significant differences by cannabis group in global or regional brain volumes, cortical thickness, or gray matter density, and no significant group by age interactions were found. Follow-up analyses indicated that values of structural neuroimaging measures by cannabis group were similar across regions, and any differences among groups were likely of a small magnitude. In sum, structural brain metrics were similar among adolescent and young adult cannabis users and non-users. Our data converge with prior large-scale studies suggesting small or limited associations between cannabis use and structural brain measures in youth. Detailed studies of vulnerability to structural brain alterations and longitudinal studies examining long-term risk are indicated.

## INTRODUCTION

Cannabis is the most popular illicit drug worldwide, with approximately 183 million individuals reporting use in the past year [1]. In 2016, 35.6% of US youth in 12^th^ grade reported cannabis use, with 6% reporting daily or almost daily use [2]. Concurrent with these trends, the perceived harms of cannabis have decreased while its societal acceptance has increased [3]. Thus, associations between cannabis use and health outcomes are of great public health importance.

One proposed risk of chronic cannabis use is alterations in brain structure, especially in adolescents and young adults. Substantial neurodevelopment occurs from adolescence to the mid-20s, including increased myelination, accelerated synaptic pruning, decreased gray matter volume, increased white matter volume, and maturation of prefrontal regions and associated neural circuitry [4, 5]. The endocannabinoid system, including CB_1_ receptors involved in the psychoactive response to cannabis, has been implicated in such neurodevelopmental changes [6]. Thus, there are increasing concerns whether cannabis use during adolescence may disrupt normative trajectories of brain development.

Recent systematic reviews have concluded that frequent cannabis use in adolescence and early adulthood is associated with abnormalities in brain structure [7, 8]. However, despite accumulating research, consistent findings in this area have remained elusive. Generally, studies have focused on brain regions with a high density of CB_1_ receptors [9], including subcortical structures such as the basal ganglia, hippocampus, and amygdala, as well as the cerebellum, cingulate cortex, and prefrontal cortex. Several studies have reported associations between frequent cannabis use in adolescents and young adults and reductions in hippocampal volumes [10–12]. However, other studies do not replicate these reductions [13–17], including one longitudinal study [18]. Similarly, orbitofrontal cortex volumes have been examined, with mixed results [19, 20]. While three studies have found larger volumes of cerebellar structures in adolescent frequent cannabis users [13, 21, 22], three also report equivocal findings or decreases in cerebellar volumes [17, 23, 24]. Frequent cannabis users have also been shown to have thinner prefrontal cortex [15, 25], although several studies have not replicated these findings [14, 26, 27]. Finally, inconsistent results are apparent in amygdala, striatum, and cingulate cortex [10, 13, 15], despite their high density of CB_1_ receptors.

These inconsistencies may be due to differences in sampling, design, and analytic techniques. For example, small sample sizes have plagued neuroscience until recently, resulting in suboptimal statistical power and reducing the precision and reliability of findings [28]. Brain imaging studies of cannabis users are not exempt from this critique. In addition, there is heterogeneity in how studies define “problematic” cannabis use, varying from a diagnosis of cannabis use disorder, criteria for number of lifetime uses, or recent frequency of use. Another factor contributing to inconsistency may be the application of numerous neuroimaging data processing pipelines and analytic techniques across the literature combined with some opacity about how such differences may affect results [29, 30]. For example, many studies apply a region of interest (ROI) approach, which may bias results (and systematic reviews) without correction for multiple comparisons, especially if ROIs were not selected *a priori*.

In the current study, we leverage a large sample of adolescents and young adults ascertained in the Philadelphia Neurodevelopmental Cohort [PNC; 31] to examine structural brain differences related to levels of cannabis use. Replicating prior work with larger samples is increasingly recognized as essential, especially with rapid policy shifts regarding cannabis. However, we also advance prior research in several ways. First, we examine both frequent and occasional cannabis users, with use criteria informed by prior research. Since the majority of cannabis users are not daily or even almost daily users, this group of occasional users is of scientific interest both as an understudied but germane sample and to allow for examination of dose-response effects. Second, to reduce variability from analytic techniques, we apply robust quality control procedures [32] and implement two well-validated automated image analysis tools. We also add parameters informative for understanding brain development, such as gray matter density [GMD; 33]. Finally, we use a community-based sample not enriched for substance use or psychopathology, enhancing generalizability.

## MATERIALS AND METHODS

### Participants

The PNC is a single-site, community-based study of 9,498 youths aged 8-21. Data from the initial cross-sectional sample, reported here, was collected from November 2009 to December 2011. For extensive details on the PNC, refer to [31, 34]. Briefly, participants were drawn from a pool of 50,293 youths recruited and genotyped by the Center for Applied Genomics at the Children’s Hospital of Philadelphia (CHOP). Participants were selected at random after stratification by sex, age, and race. Importantly, selection was not based on psychiatric or substance use symptoms. PNC inclusion criteria included living in the tristate area of Pennsylvania, New Jersey, and Delaware; proficiency in English; and ability to provide informed consent/assent. Exclusion criteria were significant developmental delays or physical conditions that would interfere with participation in neurocognitive and psychiatric assessments (e.g., pervasive developmental disorders). Criteria were intentionally broad in order to enhance generalizability. Participants who received neuroimaging were excluded for standard MRI contraindications. Participants and their guardians (for participants under 18) provided written informed consent/assent. Institutional Review Boards at University of Pennsylvania and CHOP approved the protocol. See Figure S1 for a flowchart of sample construction.

### Substance Use Assessment

Details of the PNC substance use assessment have been reported in detail [35]. Briefly, most participants received an abbreviated version of the Minnesota Center for Twin and Family Research self-report substance use assessment [36], which was privately self-administered on a laptop computer. The measure assessed lifetime use of several substances, and for cannabis and alcohol, additional questions gathered details about age at first use, age at first daily use, frequency of past year use, and methods of access. In the initial phase of the project, prior to implementing the self-report measure, an assessor-administered version of the Kiddie-Schedule for Affective Disorders and Schizophrenia (K-SADS) substance screener was administered. Since this screener did not query frequency of use, here we only include the subset of participants who denied cannabis use from those who were only administered the K-SADS screener. This measure was subsequently replaced to accommodate high participant volume and reduce social desirability biases. To further reduce social desirability biases, participants were assessed separately from collaterals (e.g., parents) and informed that information reported would be kept confidential except as legally required. Participants endorsing use of fake drug items (e.g., “cadrines”) were excluded from analyses (*n*=8), similar to previous work [35].

Analyses were conducted in participants between 14 and 21 years old (*n*=781) given limited cannabis use by those under 14 [see 35]. Informed by prior work from our group [35] and others [37, 38], we divided cannabis users into Frequent Users (≥ “3-4 times per week”; *n*=38) and Occasional Users (≤ “1-2 times per week”; *n*=109). Information regarding abstinence and urine toxicology were not acquired. To examine associations with cannabis from cumulative recent use, as opposed to remote use, we only examined cannabis users who endorsed use over the past year, removing 62 participants from analysis.

Additional sensitivity analyses evaluated alcohol consumption. A covariate was created by summing the *z*-scores from five questions addressing frequency of drinking episodes, number of drinks per episode, maximum number consumed in previous drinking episodes, and frequency of intoxication, all within the last 12 months. Given the skewness of these data, summed *z*-scores were scaled between 1 and 2 and logarithmically transformed. This covariate was included in supplementary analyses. Alcohol data were not available for *n*=35 participants.

### Psychopathology Assessment

As described previously [31], a computerized version of the K-SADS collected information on symptoms, duration, distress, and impairment for lifetime mental health symptoms. Using these item-level data, empirically-derived psychopathology factor scores were generated from a bi-factor confirmatory model, parsing psychopathology into a general factor, representing the overall burden of psychopathology, and four orthogonal symptom dimensions representing anxious-misery (mood and anxiety), fear (phobias), behavioral/externalizing, and psychosis-spectrum symptoms [39]. To control for the influence of psychopathology, given its relevance for brain morphometry in previous studies [40], overall psychopathology was included as a covariate in all analyses.

### Neuroimaging Acquisition

A subset of the PNC (*n*=1601) received structural MRI, as previously reported [34]. Imaging data was acquired on a single MRI scanner (Siemens 3T TIM Trio, Erlangen, Germany; 32-channel head coil) using the same sequences for all participants. A magnetization-prepared, rapid acquisition gradient-echo (MPRAGE) T1-weighted image was acquired with the following parameters: TR 1810 ms; TE 3.51 ms; FOV 180×240 mm; matrix 192×256; 160 slices; slice thickness/gap 1/0 mm; TI 1100 ms; flip angle 9 degrees; effective voxel resolution of 0.93×0.93×1.00 mm; total acquisition time 3:28 minutes. As detailed previously [32], three highly trained image analysts independently assessed structural image quality. Images with gross artifacts were excluded from analyses. To control for the confounding influence of image quality, we also included the average quality rating as a covariate in all models.

### Advanced Normalization Tools (ANTs) Structural Image Processing

We evaluated multiple measures including cortical thickness (CT), volume, and GMD. CT was quantified using tools in ANTs [41]. To avoid registration bias and maximize sensitivity to detect regional effects that can be impacted by registration error, a custom adolescent template and tissue priors were created using data from 140 PNC participants, balanced for age and sex. Structural images were processed and registered to this custom template using the ANTs cortical thickness pipeline [42], which includes brain extraction, N4 bias field correction [41], Atropos tissue segmentation [43], SyN diffeomorphic registration [44], and diffeomorphic registration-based CT (DiReCT) estimation in volumetric space [45]. Regional CT was taken by averaging CT estimates over anatomically defined regions, as defined below.

To parcellate the brain into anatomically-defined regions, we used an advanced multi-atlas labeling approach. Specifically, 24 young adult T1-weighted volumes from the OASIS data set [46], manually labeled by Neuromorphometrics, Inc., were registered to each subject’s T1-weighted volume using SyN diffeomorphic registration [44]. Label sets were synthesized into a final parcellation via joint label fusion [47]. Volume was determined for each parcel using the intersection between the parcel created and a prior-driven gray matter cortical segmentation from the ANTs cortical thickness pipeline to increase tissue specificity.

Finally, GMD was calculated via Atropos [48], with an iterative segmentation procedure initialized using 3-class K-means segmentation. This procedure produces both a discrete 3-class hard segmentation and a probabilistic GMD map (soft segmentation) for each subject. GMD was calculated within the intersection of this 3-class segmentation and the subject’s volumetric parcellation [33]. Importantly, this method is distinct from methods in most prior studies that use GMD interchangeably with voxel based morphometry analyses [e.g., 49], which display relationships closer to volumetric analyses than to density parameters [33].

### FreeSurfer Image Processing

Cortical reconstruction of the T1 image was performed for all subjects using FreeSurfer version 5.3 [50]. See Supplementary Methods for detailed procedures.

### Statistical Analyses

Differences among demographic and psychopathology variables were explored using ANOVAs, chi-square tests, and pairwise t-tests. Differences across groups on imaging variables were explored using ANOVAs and Kruskall-Wallis tests, while controlling for race, sex, overall psychopathology, average quality rating, and nonlinear effects of age. Nonlinearities were handled using generalized additive models (GAMs) with penalized splines as implemented in the `mgcv` package in R. Interactions between groups and age, quadratic, and cubic expansions of age were explored in a similar framework. Group differences in brain measures were examined across levels of anatomical specificity, including global (mean GMD, total brain volume [TBV], cortical thickness), lobular (frontal, temporal, occipital, parietal, insular, limbic), and regional values. To explore significant omnibus ANOVAs, pairwise relationships were explored using t-tests. False discovery rate (FDR) correction was used to account for multiple comparisons throughout. Analyses were conducted in R version 3.3.

Follow-up nonparametric data-driven analyses were run to further probe for pairwise group differences while limiting the potential influence of non-normal distributions or outliers. Mean differences were estimated across 10,000 bootstrap folds as implemented by the `boot` package in R. Studentized 95% confidence intervals were then obtained. To account for multiple comparisons, *p*-values were obtained for every confidence interval as previously detailed [51], and FDR correction was applied.

As described below, these analyses did not reveal significant effects. Since non-significant tests do not necessarily support null results, we performed follow-up equivalence tests to examine whether the presence of effects of a particular magnitude could be statistically rejected, allowing for greater specificity in defining the magnitude of potential group differences. Two one-sided *t*-tests (TOSTs) evaluated equivalence between each pairwise comparison [52] as implemented in the ‘equivalence’ package in R. TOSTs require an upper and lower bound effect size. Due to increased sample sizes required to conduct TOSTs [52], effect sizes conventionally considered to be medium magnitude were first examined, setting our equivalence boundary at *d* = -.5 and .5, respectively. A follow-up analysis used an effect size boundary from *d* = -.3 to .3 to compare Occasional and Non-Users (Frequent User comparisons were not conducted at this boundary due to limited power). Two composite *t*-tests were run, one probing larger and the other smaller than the prespecified boundaries. In these tests, the null hypothesis is non-equivalence, or the presence of an effect.

## RESULTS

### Demographic and Substance Use Data

Both cannabis groups were older and included a higher proportion of males than Non-Users, with Frequent Users having a higher proportion of males than Occasional Users (Table 1). Groups were similar in racial composition. Compared to Non-Users, Frequent Users evidenced lower estimated IQs as assessed by the Reading Subtest of the Wide Range Achievement Test-4 [53], although all groups had estimated IQs within the average range. Frequent Users started using cannabis at younger ages than Occasional Users, and both user groups reported more frequent alcohol use and higher overall psychopathology than Non-Users.

**Table 1.**
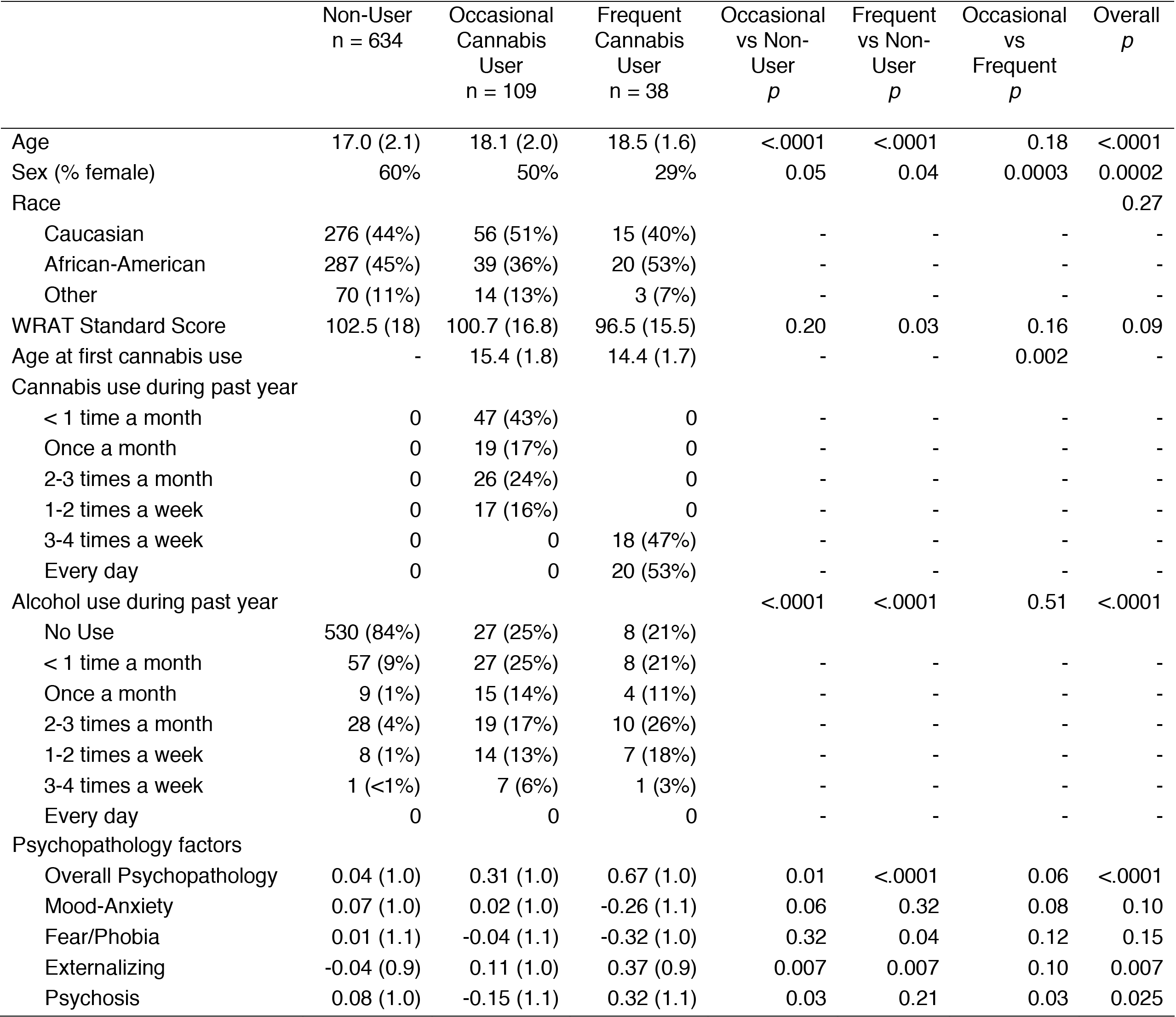
Demographic and substance use characteristics of the sample, by cannabis use group

|  | Non-User<br>n = 634 | Occasional<br>Cannabis<br>User<br>n = 109 | Frequent<br>Cannabis<br>User<br>n = 38 | Occasional<br>vs Non-User<br>*p* | Frequent<br>vs Non-User<br>*p* | Occasional<br>vs<br>Frequent<br>*p* | Overall<br>*p* |
| --- | --- | --- | --- | --- | --- | --- | --- |
| Age | 17.0 (2.1) | 18.1 (2.0) | 18.5 (1.6) | <.0001 | <.0001 | 0.18 | <.0001 |
| Sex (% female) | 60% | 50% | 29% | 0.05 | 0.04 | 0.0003 | 0.0002 |
| Race |  |  |  |  |  |  | 0.27 |
| Caucasian | 276 (44%) | 56 (51%) | 15 (40%) | - | - | - | - |
| African-American | 287 (45%) | 39 (36%) | 20 (53%) | - | - | - | - |
| Other | 70 (11%) | 14 (13%) | 3 (7%) | - | - | - | - |
| WRAT Standard Score | 102.5 (18) | 100.7 (16.8) | 96.5 (15.5) | 0.20 | 0.03 | 0.16 | 0.09 |
| Age at first cannabis use | - | 15.4 (1.8) | 14.4 (1.7) | - | - | 0.002 | - |
| Cannabis use during past year |  |  |  |  |  |  |  |
| < 1 time a month | 0 | 47 (43%) | 0 | - | - | - | - |
| Once a month | 0 | 19 (17%) | 0 | - | - | - | - |
| 2-3 times a month | 0 | 26 (24%) | 0 | - | - | - | - |
| 1-2 times a week | 0 | 17 (16%) | 0 | - | - | - | - |
| 3-4 times a week | 0 | 0 | 18 (47%) | - | - | - | - |
| Every day | 0 | 0 | 20 (53%) | - | - | - | - |
| Alcohol use during past year |  |  |  | <.0001 | <.0001 | 0.51 | <.0001 |
| No Use | 530 (84%) | 27 (25%) | 8 (21%) |  |  |  |  |
| < 1 time a month | 57 (9%) | 27 (25%) | 8 (21%) | - | - | - | - |
| Once a month | 9 (1%) | 15 (14%) | 4 (11%) | - | - | - | - |
| 2-3 times a month | 28 (4%) | 19 (17%) | 10 (26%) | - | - | - | - |
| 1-2 times a week | 8 (1%) | 14 (13%) | 7 (18%) | - | - | - | - |
| 3-4 times a week | 1 (<1%) | 7 (6%) | 1 (3%) | - | - | - | - |
| Every day | 0 | 0 | 0 | - | - | - | - |
| Psychopathology factors |  |  |  |  |  |  |  |
| Overall Psychopathology | 0.04 (1.0) | 0.31 (1.0) | 0.67 (1.0) | 0.01 | <.0001 | 0.06 | <.0001 |
| Mood-Anxiety | 0.07 (1.0) | 0.02 (1.0) | -0.26 (1.1) | 0.06 | 0.32 | 0.08 | 0.10 |
| Fear/Phobia | 0.01 (1.1) | -0.04 (1.1) | -0.32 (1.0) | 0.32 | 0.04 | 0.12 | 0.15 |
| Externalizing | -0.04 (0.9) | 0.11 (1.0) | 0.37 (0.9) | 0.007 | 0.007 | 0.10 | 0.007 |
| Psychosis | 0.08 (1.0) | -0.15 (1.1) | 0.32 (1.1) | 0.03 | 0.21 | 0.03 | 0.025 |

### Structural Imaging Analyses

Group differences were non-significant across ranges of anatomical specificity. There were no significant group differences in any global metric (TBV, total white matter volume, total gray matter volume, mean CT, mean GMD) from the ANTs processing pipeline (see Figures 1 & 2) or the FreeSurfer pipeline (see Figures S1 & S2 and Supplementary Results). Consistent, non-significant results were found using Kruskall-Wallis tests. At the lobular level, there were no significant group differences in volume or GMD. For cortical thickness (using ANTs), two lobular regions were nominally significant: the left frontal lobe (*F*=4.60, *p*=0.01) and the left parietal lobe (*F*=3.29, *p*=0.04) (Figure 1). After an FDR threshold of Q=.05 was applied, these omnibus tests were no longer significant.

**Figure 1.**
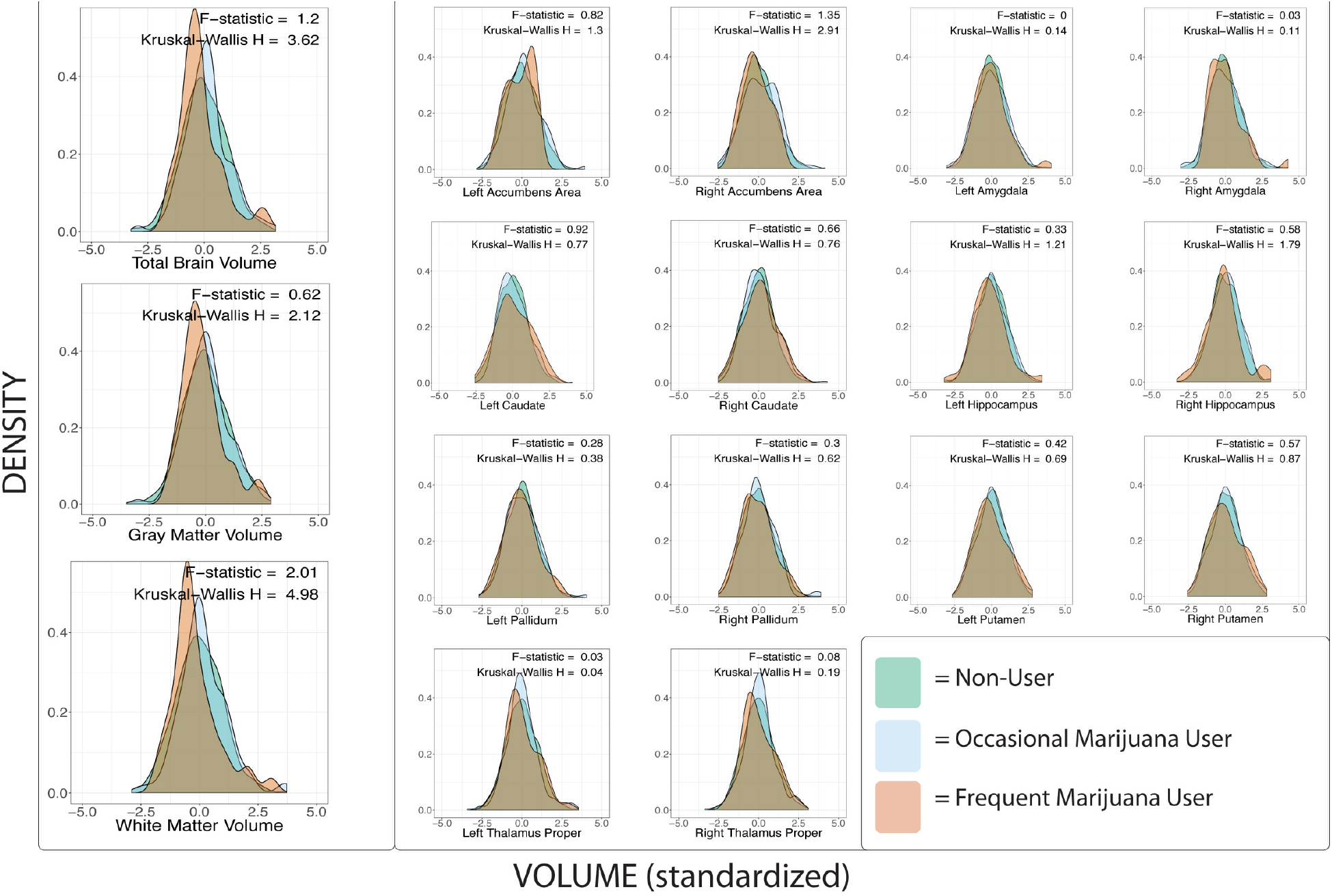
Density plots across the three groups of interest displaying standardized cortical thickness values using the Advanced Normalization Tools (ANTs) cortical thickness pipeline. Prior to plotting nonlinear effects of age, sex, mean manual rating quality, psychopathology, and race effects were removed from the values. On the left, mean cortical thickness is plotted, on the right lobular specific values are plotted.

**Figure 2.**
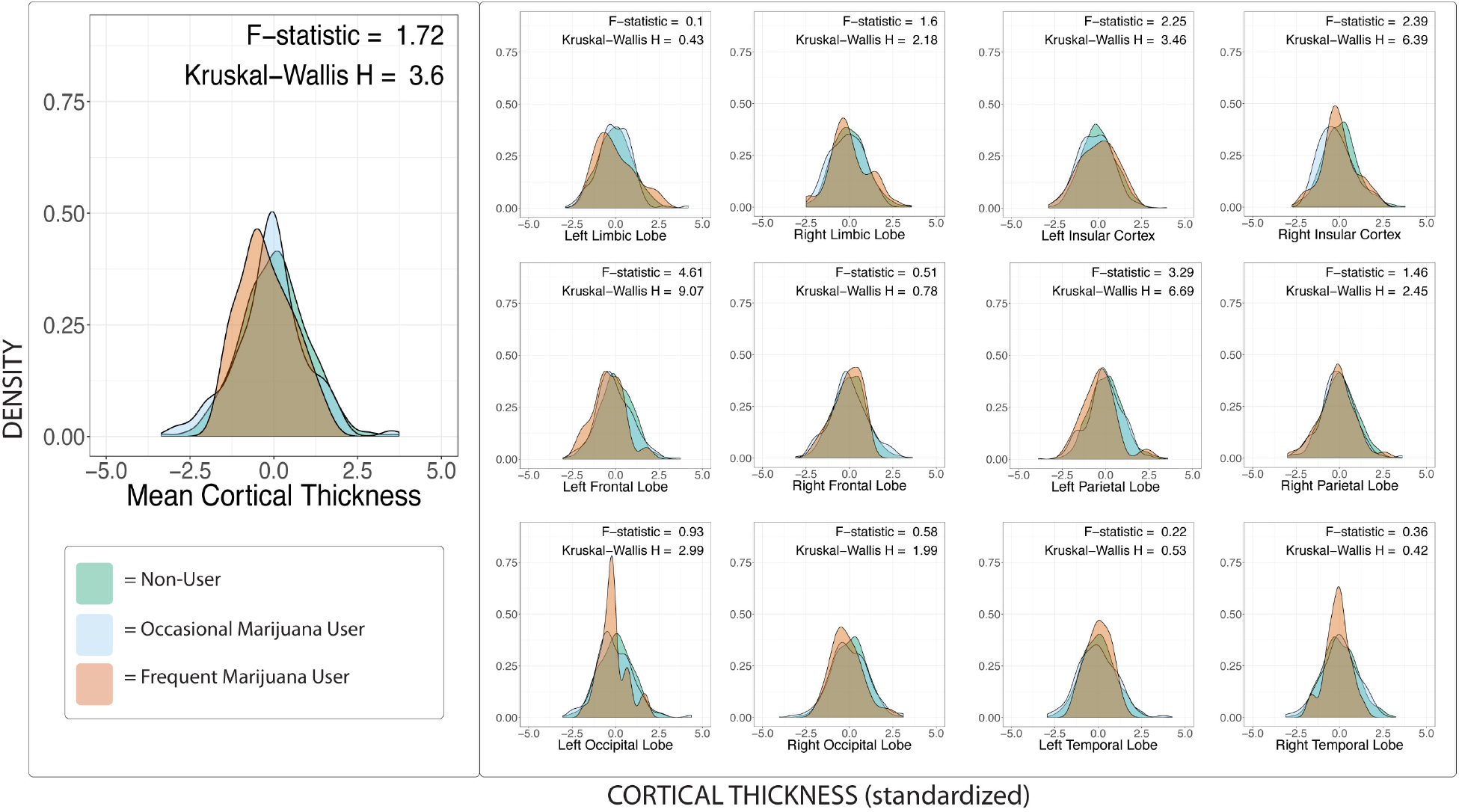
Density plots across the three groups of interest displaying standardized volume values using Advanced Normalization Tools (ANTs). Prior to plotting, nonlinear effects of age, sex, mean manual rating quality, psychopathology, and race effects were removed from the values. On the left, global metrics are plotted, while subcortical regions are plotted on the right.

For regional differences, four regions of interest yielded uncorrected significant results in volume, including the left posterior cingulate gyrus, right superior temporal gyrus, and bilateral cerebellar white matter, none of which remained significant after FDR correction (see Table S1). Cortical thickness analyses found 11 uncorrected significant regions of interest, while none remained significant after FDR correction (see Table S1). There were no significant group differences in GMD. Effect sizes across regions in cortical thickness are presented in Figures S4-S6 to discern overall trends in the data, and brain slices mapping regional *F* and *t* values and analytic code are available online (https://adrose.github.io/nullef/index.html).

Analyses of age by group interactions also displayed non-significant effects. Across all 396 comparisons, no test yielded a significant interaction.

### Supplemental Analyses

When analyses were re-run including alcohol use in the model, findings were convergent with prior results, such that there were non-significant differences among groups across all levels of anatomical specificity.

### Bootstrap Analyses

We followed these results with nonparametric, data-driven bootstrap analyses. Largely, results remained consistent (Figure 3 shows global results), suggesting minimal group differences in cortical thickness, volume, or gray matter density at the global, lobular, or regional levels. However, one pairwise difference between Non-Users and Frequent Users in left frontal lobe CT remained significant after FDR correction (95% CI=[.14,.75], *z*=2.9, *p*=.041), indicating greater CT in Non-Users.

**Figure 3.**
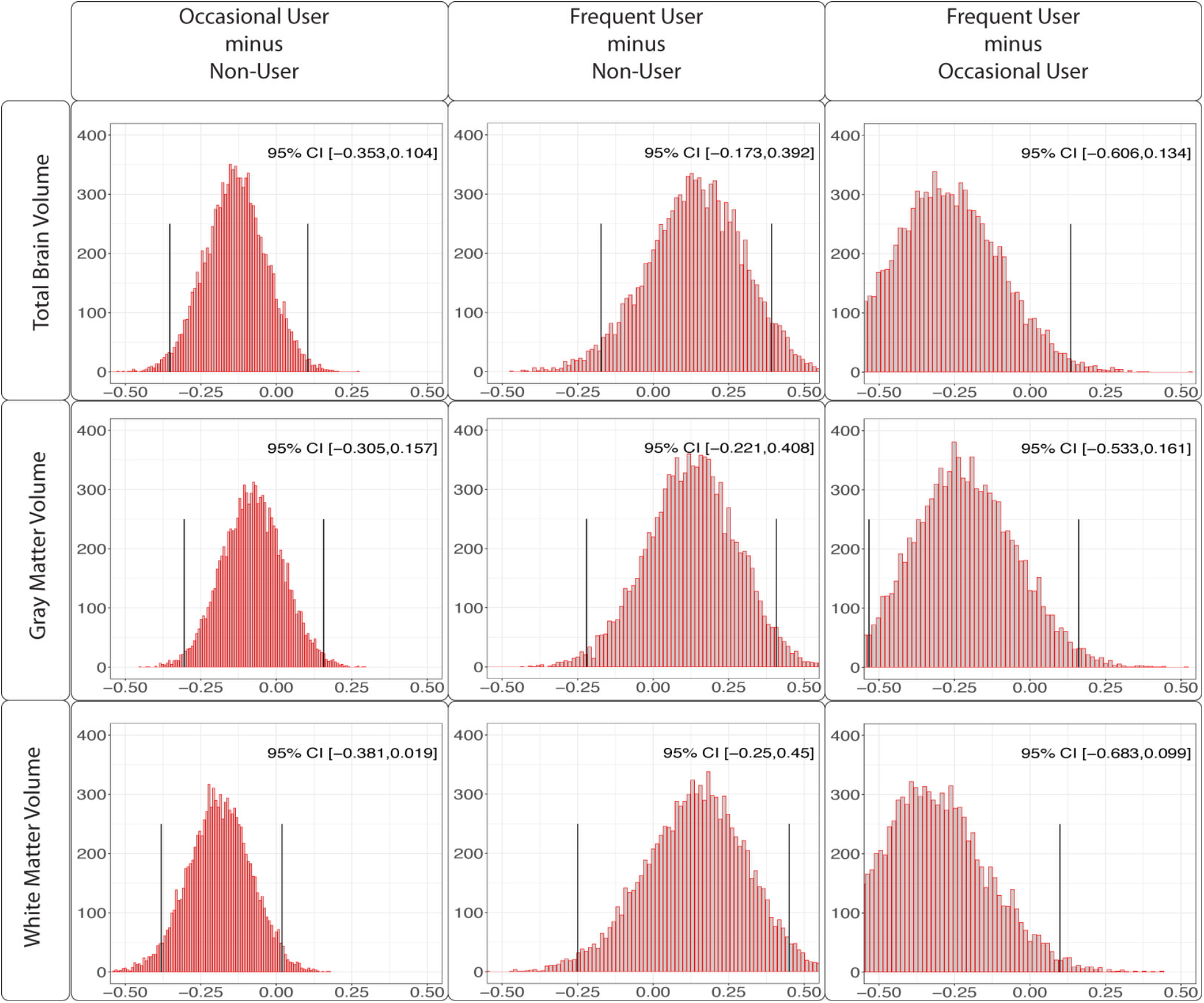
Histograms of mean differences across 10,000 bootstrap folds are displayed above for total brain volume, total gray matter volume, and total white matter volume. P values were calculated by counting the number of mean differences across the 10,000 folds that had an absolute difference greater than .3. Vertical lines display the average of the mean differences +/- the standard error.

### Equivalence Testing

Finally, we implemented equivalence testing to examine the inconsistent, relatively small effects reported above. At all levels of anatomical specificity, TOSTS remained significant at an FDR threshold of Q=.05 for all contrasts (e.g., Frequent vs. Non-Users), indicating that any differences across groups in brain structural measures were between *d* = .5 and -.5. Follow-up TOSTs were run limiting the magnitude of the effects to *d* = +/-.3 to compare the Non-Users and Occasional Users, and all TOSTS remained significant, suggesting equivalence.

## DISCUSSION

In the current study, we used a systematic approach to examine whether occasional or frequent cannabis use was associated with alterations in measures of brain structure in a large, community-based, single-scanner sample of adolescents and young adults. Using rigorous quality control procedures and two well-validated analysis software programs, we did not replicate previously reported structural differences in cannabis users, as we found few differences in brain structural measures associated with either occasional or frequent cannabis use in adolescence. Although the distribution of effect sizes may suggest slightly lower levels of cortical thickness in prefrontal regions in frequent users compared to both occasional users and non-users, none of these differences reached statistical significance after controlling for multiple comparisons. We also statistically evaluated our non-significant results and provided support for the absence of medium or greater magnitude effects across groups. In addition, we did not find significant interactions between cannabis use and age, which would have suggested increased vulnerability to cannabis use at younger ages.

Several studies have reported significant associations between frequent cannabis use and alterations in subcortical and cerebellar volumes, cortical thickness, and surface-based morphometry in adolescents and young adults [7]. However, results have been notably variable and predominantly from studies with modest sample sizes. In the current study, we found limited evidence for structural brain differences, especially between occasional users and non-users of cannabis. Moreover, few structural neuroimaging metrics showed a coherent pattern of dose-response relationships by level of cannabis use. We also found minimal evidence of brain structural differences of a medium magnitude between frequent cannabis users and the other two groups, although the presence of smaller magnitude differences cannot be ruled out. Previous studies with small samples were likely underpowered to detect small magnitude effects, which could partially explain variability in the literature. Our results converge with data from larger samples of cannabis-using youth, which have found more limited brain structural differences associated with cannabis than smaller studies. For example, Weiland and colleagues [17] compared 50 adolescent daily users of cannabis to 50 demographically matched non-users, replicating methods from an earlier study [49], and found non-significant differences in volume, surface-based morphometry, and shape. Similarly, in a sample of 439 adolescents, Thayer and colleagues [54] found no significant associations of past month cannabis use with brain volume or measures of diffusivity after covarying for alcohol use disorder symptoms. Our results extend these studies by probing effects across levels of cannabis use and examining measures of cortical thickness and gray matter density.

Although non-significant results from null hypothesis significance testing should not be interpreted as supporting “null” findings, we performed follow-up equivalence testing to provide context for these results. These analyses suggested that differences between groups across structural imaging metrics were likely less than *d*=.5, conventionally considered to be a medium magnitude effect size, and any differences between occasional users and non-users of cannabis were likely less than *d*=.3. As an example to aid in interpreting a *d*=.5 difference, there is a 64% chance that a person picked randomly from the frequent users would have a lower value in a measure such as cortical thickness than a person picked at random from the non-users (see http://rpsychologist.com/d3/cohend/). For effect sizes smaller than *d*=.5, there would be even lower probabilities of smaller values for a person picked randomly from the frequent cannabis group.

An alternative explanation of findings is that neuroanatomical alterations may only be present in youth with: a) heavier use than observed here, and/or b) symptoms of abuse or dependence. While we cannot rule out these possibilities given our limited information about cannabis use disorders, the criteria and use patterns of our sample appear similar to those from prior adolescent studies invoked to support the presence of structural brain differences [e.g., 16, 20]. In addition, our user groups had higher levels of alcohol use and more psychopathology, reflecting expected sample characteristics. Moreover, frequent users had higher levels of these symptoms than less frequent users, although dose-response relationships in structural brain metrics were not apparent. It is also possible that structural brain alterations require more of a cumulative dose than observed here. Longitudinal research is needed to address such questions.

### Considerations for Interpretation

It is challenging to integrate our findings with the overall literature regarding cannabis and brain structure because of methodological heterogeneity. For example, some studies follow group-level analyses (sometimes without significant differences) with correlational analyses between structural metrics and variables such as age at first cannabis use or cannabis quantity; yet, whether these represent post-hoc analyses, increasing Type I error risk, is rarely discussed. Heterogeneity in findings may also reflect potential moderators of risk/vulnerability that are either unidentified to date or untested because of limited statistical power. For example, in a large sample of adolescents between the ages of 12 and 21, French and colleagues [55] found that cannabis use before age 16 was associated with slightly reduced cortical thickness, but only in males with a high polygenic risk score for schizophrenia.

Additionally, causal inferences from observational data should be undertaken cautiously. Reflecting the complexity of interpreting brain structural differences, Cheetham and colleagues [56] showed that smaller orbitofrontal cortex volumes at age 12 were predictive of cannabis initiation at age 16, suggesting that some structural brain differences could reflect pre-use risks as opposed to consequences of use. In adolescent samples, it is also challenging to determine whether group differences reflect “disruptions,” as neurodevelopment is dynamic and dependent on complex trajectories of gray and white matter development [57]. To this end, longitudinal studies have reported mixed findings regarding altered trajectories of brain structural measures in cannabis users [27, 58].

Although we did not find strong support for brain structure alterations in cannabis users, our study cannot answer whether cannabis affects brain *functioning* in adolescent-onset users. Long-term studies of brain changes are scarce, and small structural effects could nonetheless have clinical significance for brain development, cognitive functioning, or mental health. It is also important to place these results in the context of the overall cannabis literature, as they do not speak to other potential risks of use, such as propensity for addiction, motivational difficulties, or mental health disorders, including psychosis.

### Limitations

Limitations of this study include the self-report of substance use and lack of objective indicators of cannabis use. Additional biological measurements were considered but lacked feasibility with the large-scale data collection required. However, our substance use assessment was selected to minimize reporting bias and has shown good reliability greater disclosure of substance use compared to thorough diagnostic interviews [59]. Moreover, although our ascertainment methods limited selection biases, it is unknown whether our data generalize to individuals with cannabis use disorders. We also did not examine measures of white matter organization from diffusion MRI or perform shape analysis, although there are significant complexities in interpreting typically used measures (e.g., fractional anisotropy) from these techniques [60, 61]. Our data were cross-sectional, which presents a particular limitation in youth samples with protracted, complex trajectories of brain development. Absent a randomized controlled trial, longitudinal data will provide the best test of whether cannabis causes structural alterations to the brain. In addition, although we controlled for group differences in age, sex, mental health symptoms, and alcohol use, these differences could have influenced results.

### Conclusions

In a large sample of adolescents and young adults, the present study found predominantly non-significant differences in brain structural measures among cannabis non-users, occasional users of cannabis, and frequent users of cannabis. Follow-up analyses indicated that these non-significant differences were less likely due to reduced statistical power and more likely to limited or smaller magnitude effects. Results diverge from some prior studies that have reported structural differences associated with cannabis in brain volumes and cortical thickness; such differences could reflect our use of larger samples and a community ascertainment approach. Longitudinal studies are needed to determine whether adolescent cannabis use is associated with longer-term changes in brain structure.

## Supplementary Online Content

**eMethods** Supplementary Methods

**eResults** Supplementary Results

**Table S1** Results Significant at Uncorrected Levels from Cortical Thickness and Volumetric Analyses

**Figure S1** Participant Flow Chart

**Figure S2** Density plots across the three cannabis groups of interest displaying standardized volume values using FreeSurfer.

**Figure S3** Density plots across the three groups of interest displaying standardized cortical thickness values using FreeSurfer.

**Figure S4** Cohen’s *d* effect sizes comparing Non-Users to Occasional Users for cortical thickness across regions grouped by lobe.

**Figure S5** Cohen’s *d* effect sizes comparing Non-Users to Frequent Users for cortical thickness across regions grouped by lobe.

**Figure S6** Cohen’s *d* effect sizes comparing Occasional Users to Frequent Users for cortical thickness across regions grouped by lobe.

### Supplementary Methods

#### FreeSurfer Image Processing

Cortical reconstruction of the T1 image was performed for all subjects using FreeSurfer version 5.3 [1]. FreeSurfer includes registration to a template, intensity normalization, gray and white matter segmentation, and tessellation of the gray/CSF and white/gray matter boundaries [2]; cortical surfaces are inflated and normalized to a template via a spherical registration. Cortical thickness is measured as the shortest distance between the pial and the white matter tessellated surfaces [2]. The cortex was then parcellated into 40 regions [3], and cortical thickness was averaged across parcels to obtain regional cortical thickness estimates without any manual correction.

### Supplementary Results

#### ANTs Structural Imaging Results

Although no group differences were found in CT or volume after FDR correction, we conducted additional analyses to discern any weak trends in the data. Results with the largest F values are detailed here, but the directions and magnitude of differences can be found in Table S1. The left frontal lobe displayed somewhat of an ordered relationship, with Non-Users displaying the largest cortical thickness, followed by Occasional Users (though a non-significant difference from Non-Users, t=1.25, p_uncorrected_=.21), and Frequent Users displaying the smallest cortical thickness (difference from Non-User t=2.93, p_uncorrected_=0.005); the difference between Occasional and Frequent Users was not significant (t=1.84, p_uncorrected_ =.07). For volume, however, the largest F value in the right superior temporal gyrus did not display an ordered relationship. Occasional Users had the largest volume (difference from Non-User t=2.77, p_uncorrected_=.006), although Frequent Users did not show volumetric differences from Occasional Users (t=1.55, p_uncorrected_=.12) or Non-Users (t=0.12, p_uncorrected_=.90).

#### FreeSurfer Structural Imaging Results

FreeSurfer results displayed subtle differences in regional associations when compared with the results from the ANTs CT pipeline. While ANTs displayed more regional heterogeneity in prefrontal regions among the groups, FreeSurfer displayed differences across parietal, temporal, occipital, and frontal regions. Overall, 8 regions were found to have significant group differences, although none remained significant after FDR correction was applied. The largest regional heterogeneity was found in the isthmus of the right cingulate gyrus (*F*=5.98, *p*=0.002). One other regional effect was found in the right hemisphere, in the superior parietal region (*F*=3.75, *p*=0.02). The remaining six regional effects were found in the left hemisphere in the inferior parietal region (*F*=3.3, *p*=.04), inferior temporal region (*F*=3.7, *p*=.03), the lateral occipital region (*F*=3.9, *p*=.02), the middle temporal region (*F*=4.4, *p*=.01), the rostral middle frontal region (*F*=3.5,*p*=.03), and the superior frontal region (*F*=3.4, *p*=.03).

The direction of these differences remined largely consistent in 5 of the 8 significant regions, with the Non-Users displaying the largest CT values and the Frequent Users displaying the smallest mean CT (left inferior temporal region, left lateral occipital region, left rostral middle frontal region, left superior frontal region, and the right superior parietal region). Effect sizes between the Non-Users and Occasional Users remained small (Cohen’s *d* range = .02-.16, *t*- statistic range = .25-1.6*, p*-range = .81-.11) whereas differences between Non-Users and Frequent Users remained relatively consistent at medium effect sizes (Cohen’s *d* range = .34-.44*, t-statistic range= 2.4-2.9, p-range=.03-.007*). For the remaining regions, in one of them the Occasional Users displayed the greatest CT (Left inferior parietal region) with a small effect size when compared to the Non-Users (Cohen’s *d*=-.04, *t*-statistic=-.38, *p*=.7), and a moderate effect when compared to the Frequent Users (Cohen’s *d*=0.42, *t*-statistic=2.3, *p*=.03). The final region, which also displayed the largest *F*-statistic, was the right isthmus of the cingulate gyrus, where the frequent users displayed the greatest CT estimates. Here frequent users displayed small effects when compared to the Non-Users (Cohen’s *d*=-.13, *t*-statistic=-.88, *p*=.38), and medium effects when compared to the Occasional Users (Cohen’s *d*=-.54, *t*-statistic=-2.8, *p*=.006).

---

**Table S1.**
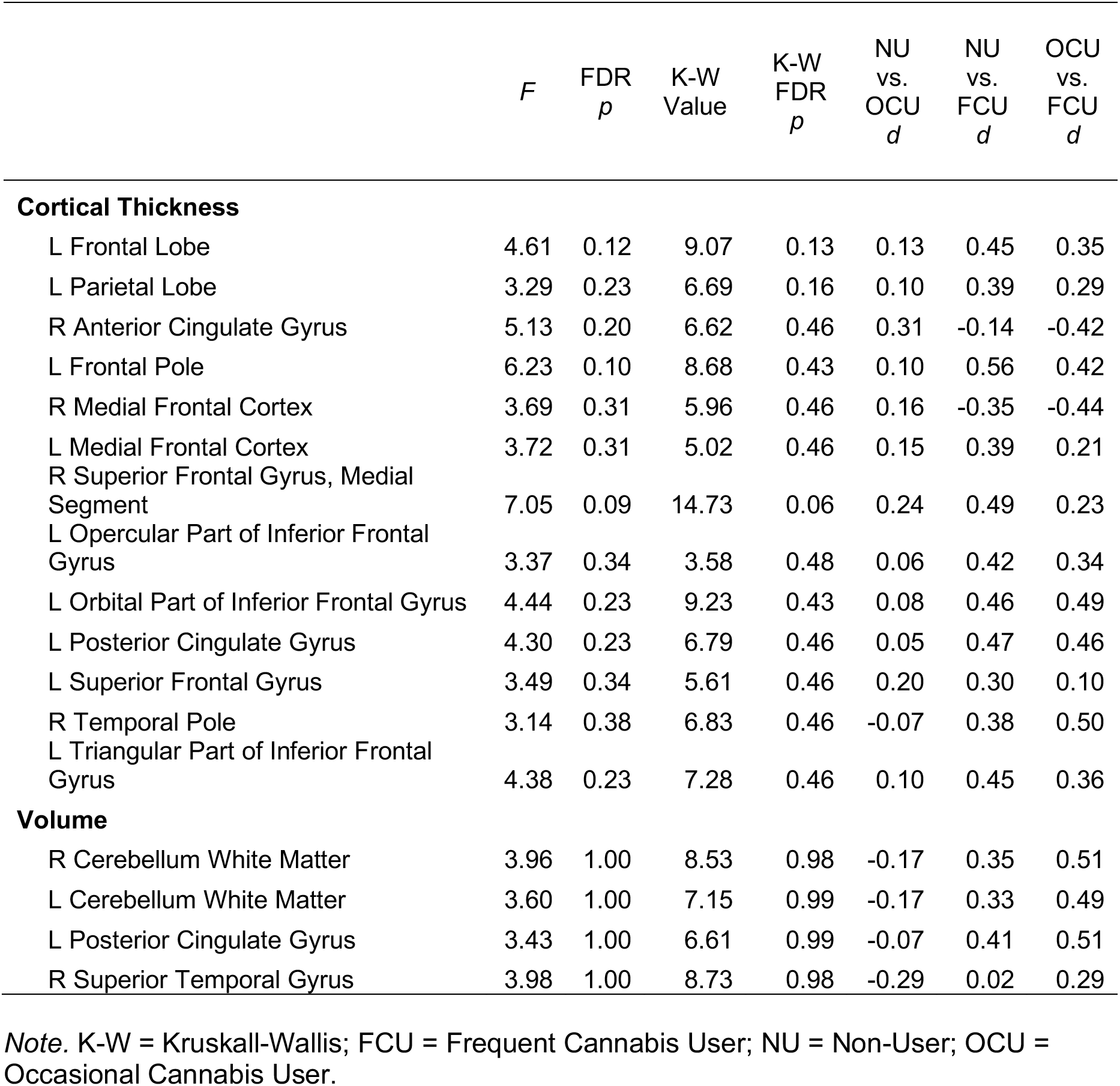
Results Significant at Uncorrected Levels from ANTs Cortical Thickness and Volumetric Analyses.

|  | *F* | FDR<br>*p* | K-W<br>Value | K-W<br>FDR<br>*p* | NU<br>vs.<br>OCU<br>*d* | NU<br>vs.<br>FCU<br>*d* | OCU<br>vs.<br>FCU<br>*d* |
| --- | --- | --- | --- | --- | --- | --- | --- |
| **Cortical Thickness** |  |  |  |  |  |  |  |
| L Frontal Lobe | 4.61 | 0.12 | 9.07 | 0.13 | 0.13 | 0.45 | 0.35 |
| L Parietal Lobe | 3.29 | 0.23 | 6.69 | 0.16 | 0.10 | 0.39 | 0.29 |
| R Anterior Cingulate Gyrus | 5.13 | 0.20 | 6.62 | 0.46 | 0.31 | -0.14 | -0.42 |
| L Frontal Pole | 6.23 | 0.10 | 8.68 | 0.43 | 0.10 | 0.56 | 0.42 |
| R Medial Frontal Cortex | 3.69 | 0.31 | 5.96 | 0.46 | 0.16 | -0.35 | -0.44 |
| L Medial Frontal Cortex | 3.72 | 0.31 | 5.02 | 0.46 | 0.15 | 0.39 | 0.21 |
| R Superior Frontal Gyrus, Medial Segment | 7.05 | 0.09 | 14.73 | 0.06 | 0.24 | 0.49 | 0.23 |
| L Opercular Part of Inferior Frontal Gyrus | 3.37 | 0.34 | 3.58 | 0.48 | 0.06 | 0.42 | 0.34 |
| L Orbital Part of Inferior Frontal Gyrus | 4.44 | 0.23 | 9.23 | 0.43 | 0.08 | 0.46 | 0.49 |
| L Posterior Cingulate Gyrus | 4.30 | 0.23 | 6.79 | 0.46 | 0.05 | 0.47 | 0.46 |
| L Superior Frontal Gyrus | 3.49 | 0.34 | 5.61 | 0.46 | 0.20 | 0.30 | 0.10 |
| R Temporal Pole | 3.14 | 0.38 | 6.83 | 0.46 | -0.07 | 0.38 | 0.50 |
| L Triangular Part of Inferior Frontal Gyrus | 4.38 | 0.23 | 7.28 | 0.46 | 0.10 | 0.45 | 0.36 |
| **Volume** |  |  |  |  |  |  |  |
| R Cerebellum White Matter | 3.96 | 1.00 | 8.53 | 0.98 | -0.17 | 0.35 | 0.51 |
| L Cerebellum White Matter | 3.60 | 1.00 | 7.15 | 0.99 | -0.17 | 0.33 | 0.49 |
| L Posterior Cingulate Gyrus | 3.43 | 1.00 | 6.61 | 0.99 | -0.07 | 0.41 | 0.51 |
| R Superior Temporal Gyrus | 3.98 | 1.00 | 8.73 | 0.98 | -0.29 | 0.02 | 0.29 |

*Note.* K-W = Kruskal-Wallis; FCU = Frequent Cannabis User; NU = Non-User; OCU = Occasional Cannabis User.

**Figure S1.**
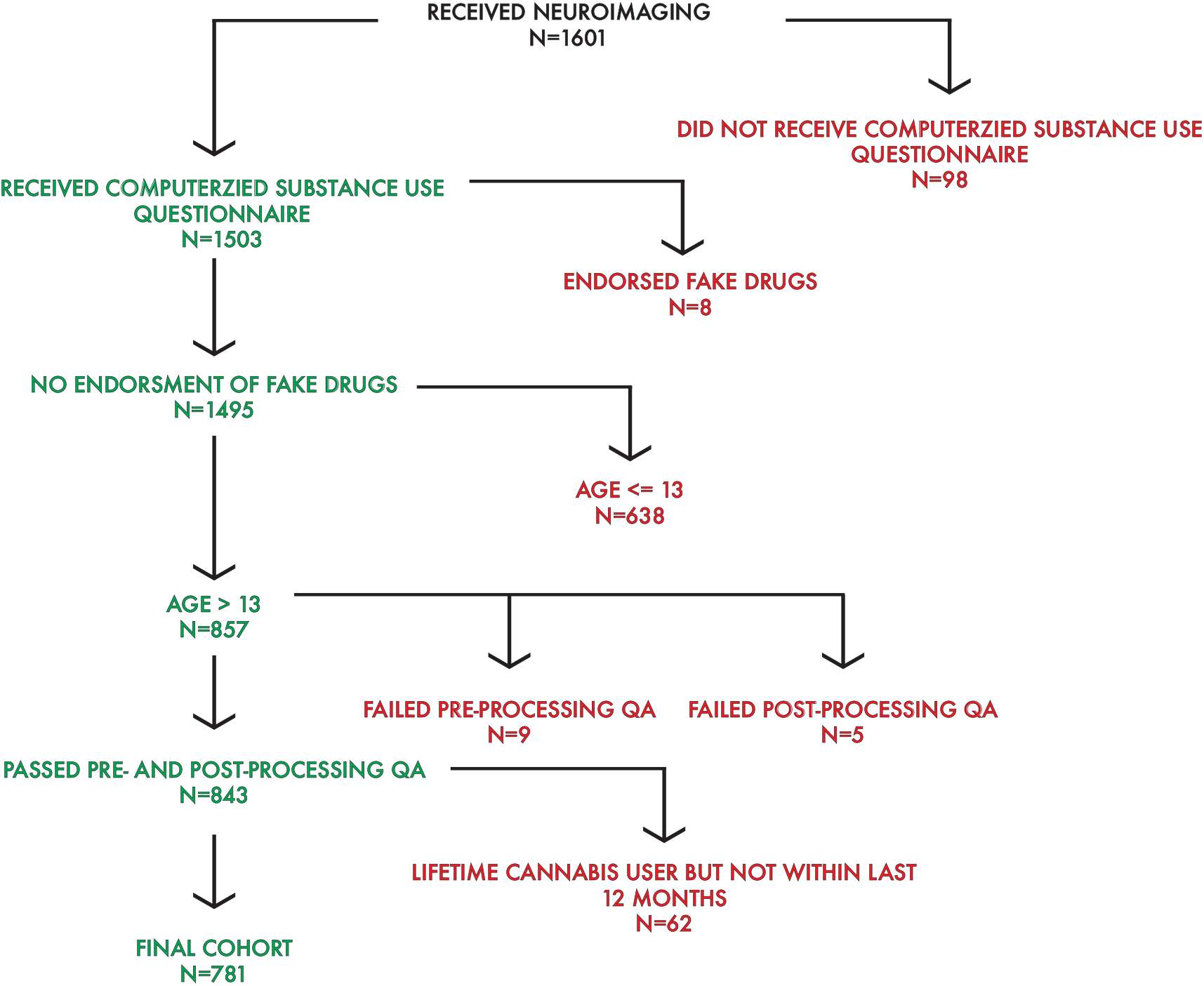
Flowchart indicating formation of the final participant sample for analysis (n=781) from the Philadelphia Neurodevelopmental Cohort.

**Figure S2.**
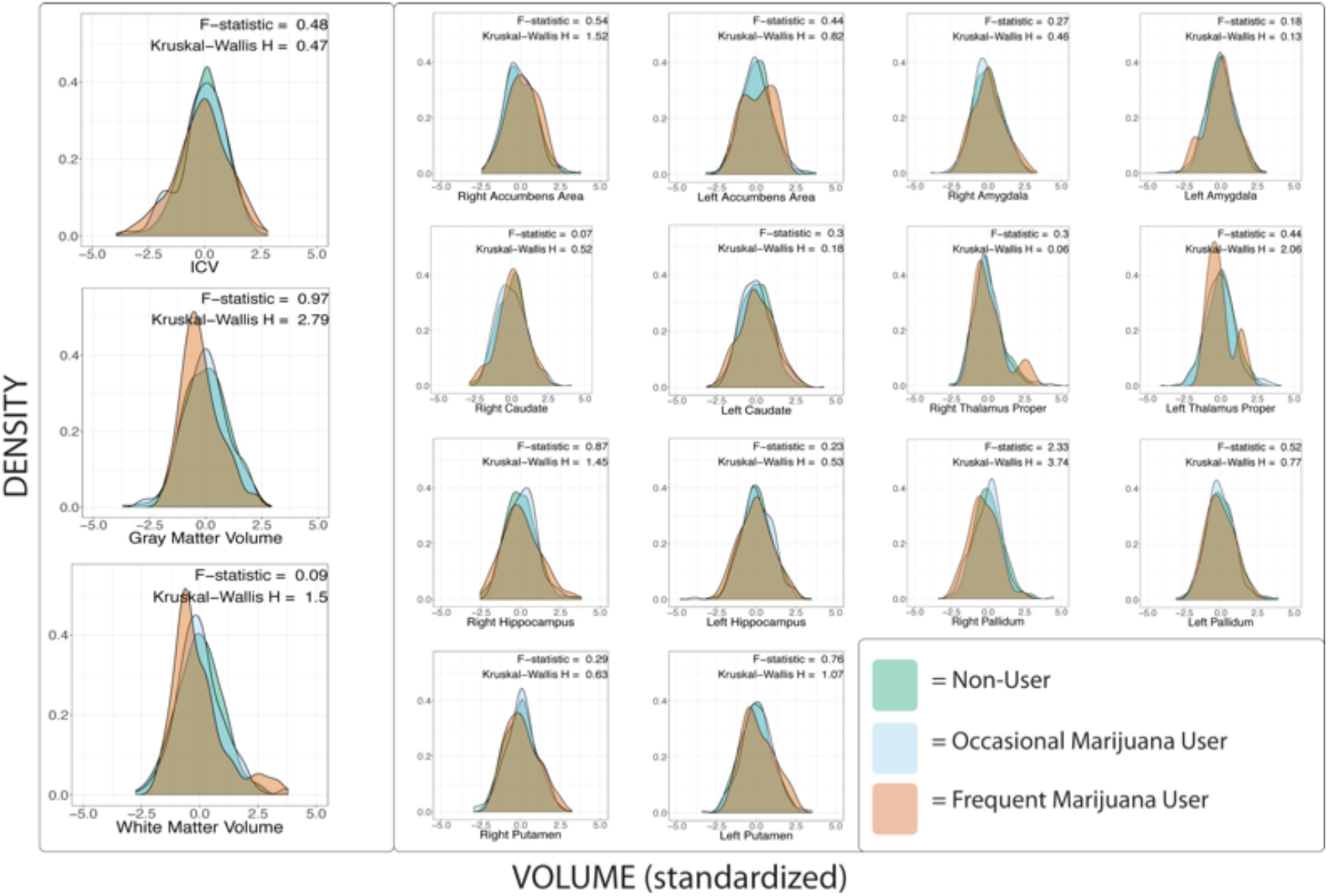
Density plots across the three cannabis groups of interest displaying standardized volume values using FreeSurfer. Prior to plotting, nonlinear effects of age, sex, mean manual rating quality, psychopathology, and race effects were removed from the values. On the left, global metrics are plotted, while subcortical regions are plotted on the right.

**Figure S3.**
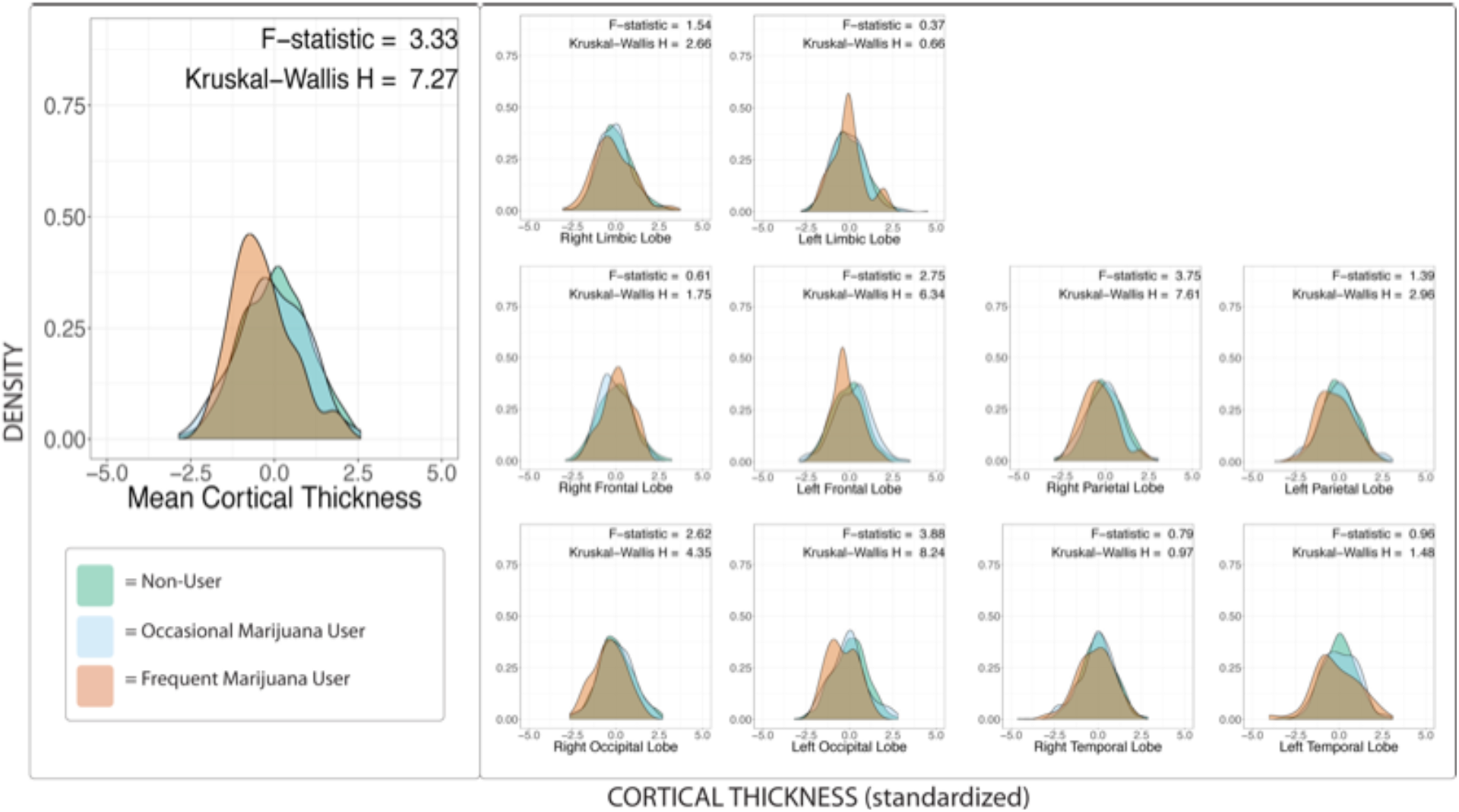
Density plots across the three groups of interest displaying standardized cortical thickness values using FreeSurfer. Prior to plotting nonlinear effects of age, sex, mean manual rating quality, psychopathology, and race effects were removed from the values. On the left, mean cortical thickness is plotted, on the right lobular specific values are plotted.

**Figure S4.**
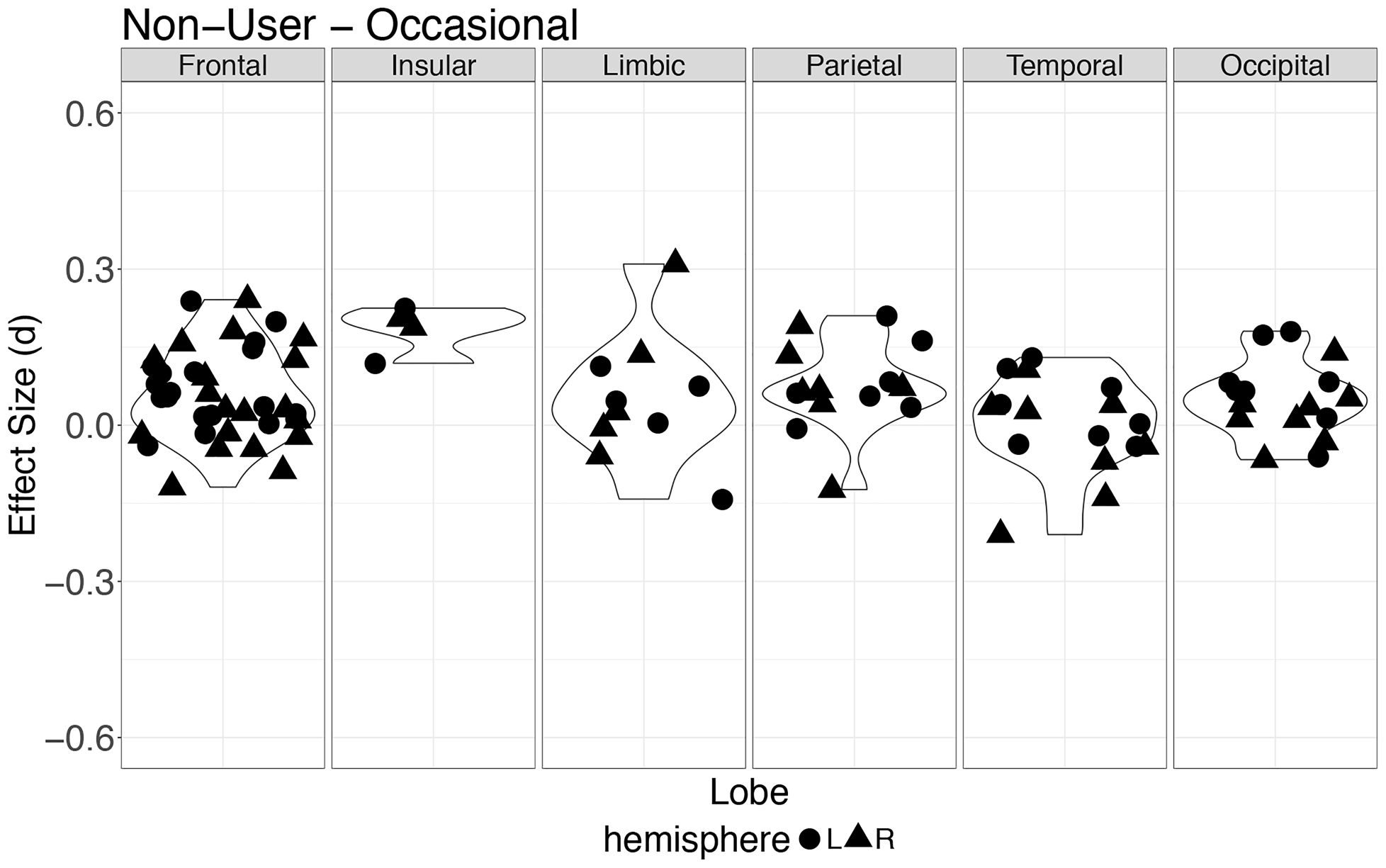
Cohen’s *d* effect sizes for cortical thickness across regions grouped by lobe, comparing Non-Users and Occasional Users. Each point represents a region within the specified lobe. Positive values reflect smaller values in the cannabis user group. Effect sizes are shown after controlling for nonlinear effects of age, sex, mean manual rating quality, psychopathology, and race.

**Figure S5.**
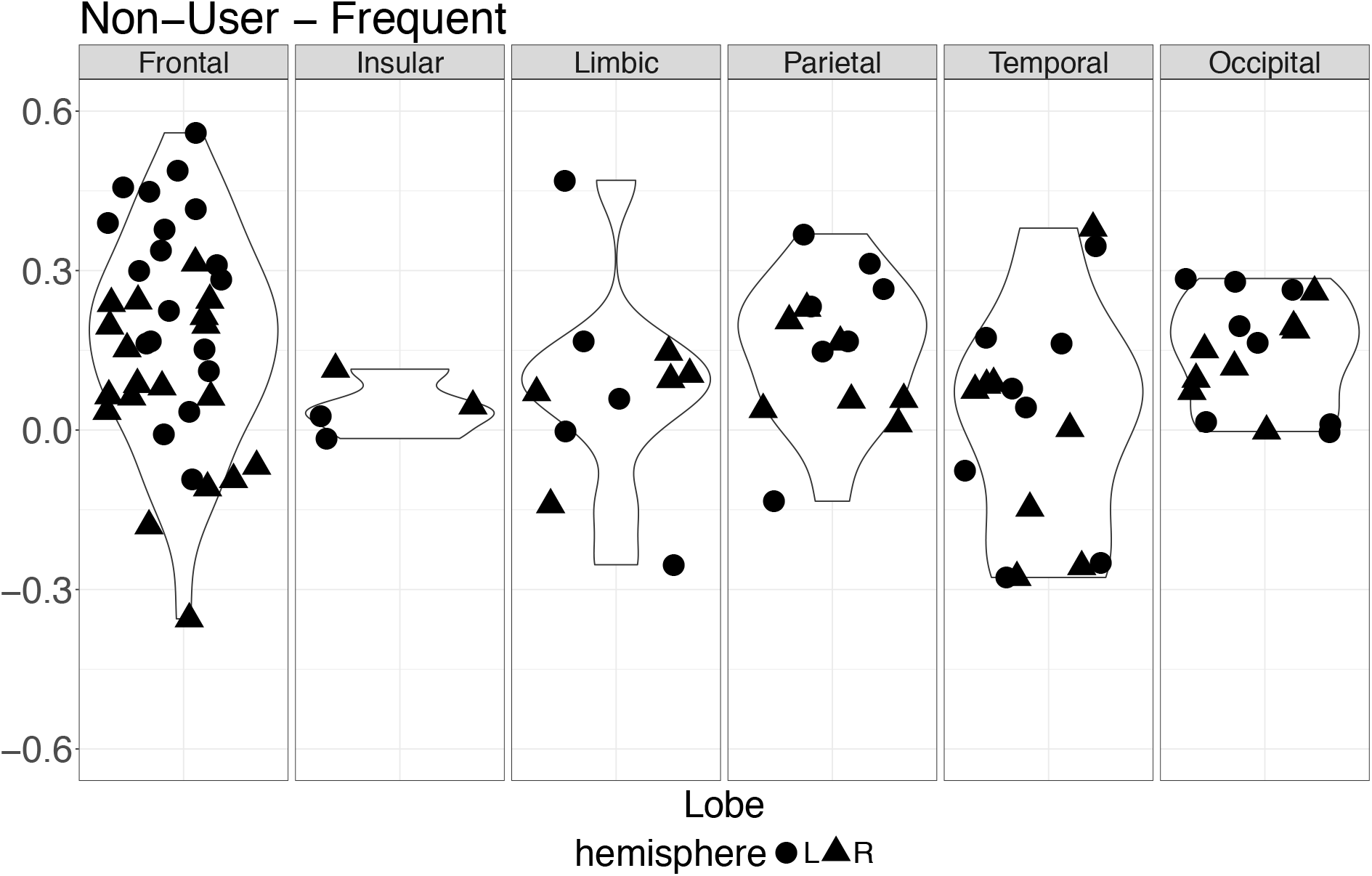
Cohen’s *d* effect sizes for cortical thickness across regions grouped by lobe, comparing Non-Users and Frequent Users. Each point represents a region within the specified lobe. Positive values reflect smaller values in the cannabis user group. Effect sizes are shown after controlling for nonlinear effects of age, sex, mean manual rating quality, psychopathology, and race.

**Figure S6.**
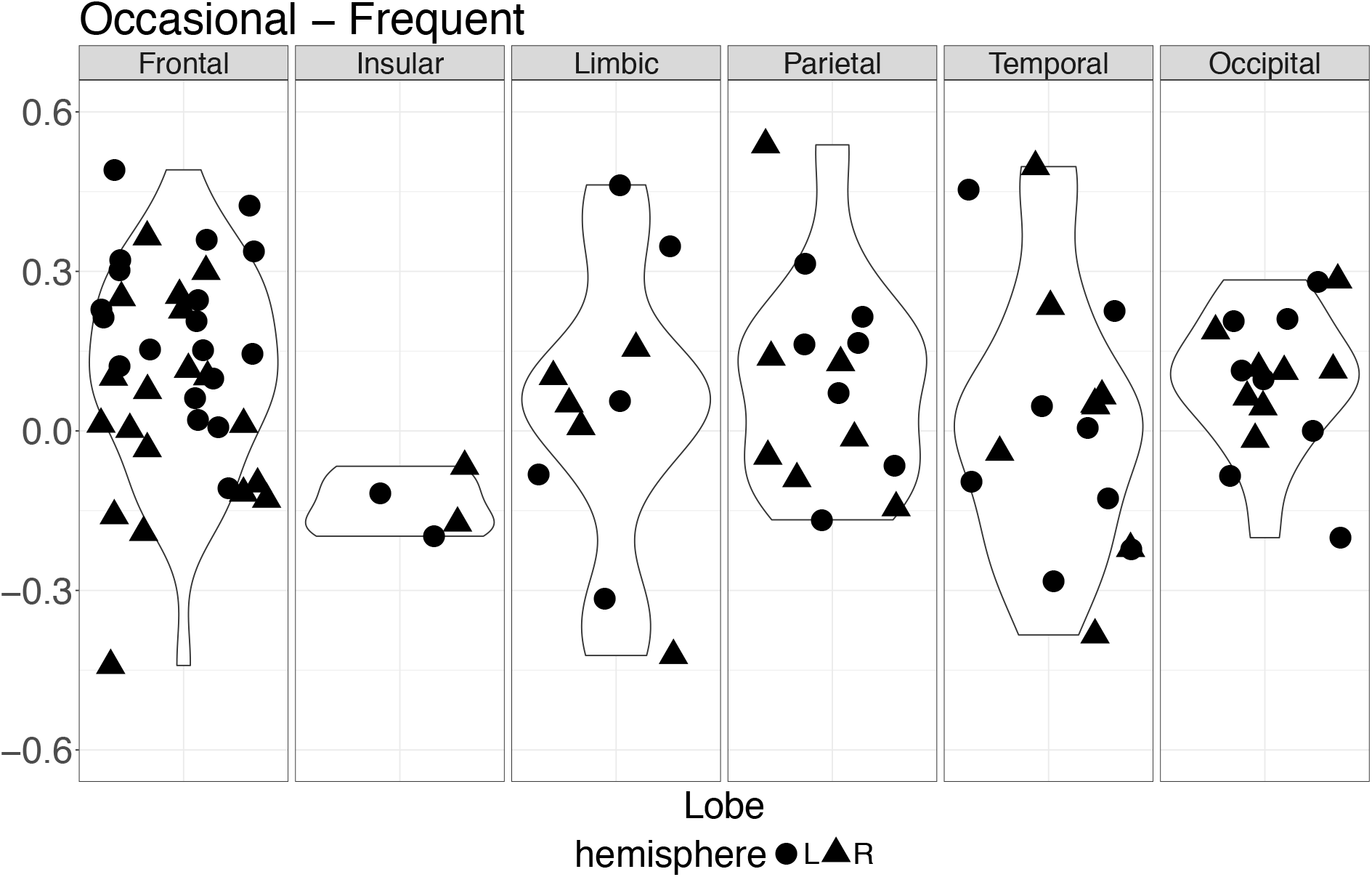
Cohen’s *d* effect sizes for cortical thickness across regions grouped by lobe, comparing Occasional Users and Frequent Users. Each point represents a region within the specified lobe. Positive values reflect smaller values in the Frequent User group. Effect sizes are shown after controlling for nonlinear effects of age, sex, mean manual rating quality, psychopathology, and race.

## REFERENCES

1. United Nations Office on Drugs and Crime. World Drug Report 2017. Vienna, Austria: United Nations Office on Drugs and Crime; 2017.

2. Johnston LD, O’Malley PM, Miech RA, Bachman JG, Schulenberg JE. Monitoring the Future national survey results on drug use, 1975-2016: Overview, key findings on adolescent drug use. Ann Arbor, MI: Institute for Social Research, The University of Michigan; 2017.

3. Substance Abuse and Mental Health Services Administration. Results from the 2013 National Survey on Drug Use and Health: Summary of National Findings. Rockville, MD: Substance Abuse and Mental Health Services Administration; 2014.

4. Mills KL, Goddings A-L, Herting MM, Meuwese R, Blakemore S-J, Crone EA, et al. Structural brain development between childhood and adulthood: Convergence across four longitudinal samples. NeuroImage. 2016;141:273–81.

5. Sowell ER, Thompson PM, Leonard CM, Welcome SE, Kan E, Toga AW. Longitudinal mapping of cortical thickness and brain growth in normal children. J Neurosci. 2004;24:8223–31.

6. Bossong MG, Niesink RJM. Adolescent brain maturation, the endogenous cannabinoid system and the neurobiology of cannabis-induced schizophrenia. Prog Neurobiol. 2010;92:370–85.

7. Batalla A, Bhattacharyya S, Yücel M, Fusar-Poli P, Crippa JA, Nogué S, et al. Structural and functional imaging studies in chronic cannabis users: a systematic review of adolescent and adult findings. PloS One. 2013;8:e55821.

8. Lorenzetti V, Solowij N, Yücel M. The Role of Cannabinoids in Neuroanatomic Alterations in Cannabis Users. Biol Psychiatry. 2016;79:e17–31.

9. Glass M, Dragunow M, Faull RL. Cannabinoid receptors in the human brain: a detailed anatomical and quantitative autoradiographic study in the fetal, neonatal and adult human brain. Neuroscience. 1997;77:299–318.

10. Ashtari M, Avants B, Cyckowski L, Cervellione KL, Roofeh D, Cook P, et al. Medial temporal structures and memory functions in adolescents with heavy cannabis use. J Psychiatr Res. 2011;45:1055–66.

11. Filbey FM, McQueeny T, Kadamangudi S, Bice C, Ketcherside A. Combined effects of marijuana and nicotine on memory performance and hippocampal volume. Behav Brain Res. 2015;293:46–53.

12. Chye Y, Suo C, Yücel M, den Ouden L, Solowij N, Lorenzetti V. Cannabis-related hippocampal volumetric abnormalities specific to subregions in dependent users. Psychopharmacology (Berl). 2017;234:2149–57.

13. Cousijn J, Wiers RW, Ridderinkhof KR, van den Brink W, Veltman DJ, Goudriaan AE. Grey matter alterations associated with cannabis use: results of a VBM study in heavy cannabis users and healthy controls. NeuroImage. 2012;59:3845–51.

14. Kumra S, Robinson P, Tambyraja R, Jensen D, Schimunek C, Houri A, et al. Parietal lobe volume deficits in adolescents with schizophrenia and adolescents with cannabis use disorders. J Am Acad Child Adolesc Psychiatry. 2012;51:171–80.

15. Mashhoon Y, Sava S, Sneider JT, Nickerson LD, Silveri MM. Cortical thinness and volume differences associated with marijuana abuse in emerging adults. Drug Alcohol Depend. 2015;155:275–83.

16. Medina KL, McQueeny T, Nagel BJ, Hanson KL, Yang TT, Tapert SF. Prefrontal cortex morphometry in abstinent adolescent marijuana users: Subtle gender effects. Addict Biol. 2009;14:457–68.

17. Weiland BJ, Thayer RE, Depue BE, Sabbineni A, Bryan AD, Hutchison KE. Daily marijuana use is not associated with brain morphometric measures in adolescents or adults. J Neurosci. 2015;35:1505–12.

18. Koenders L, Lorenzetti V, de Haan L, Suo C, Vingerhoets W, van den Brink W, et al. Longitudinal study of hippocampal volumes in heavy cannabis users. J Psychopharmacol (Oxf). 2017;31:1027–34.

19. Churchwell JC, Lopez-Larson M, Yurgelun-Todd DA. Altered frontal cortical volume and decision making in adolescent cannabis users. Front Psychol. 2010;1:225.

20. Price JS, McQueeny T, Shollenbarger S, Browning EL, Wieser J, Lisdahl KM. Effects of marijuana use on prefrontal and parietal volumes and cognition in emerging adults. Psychopharmacology (Berl). 2015;232:2939–50.

21. Battistella G, Fornari E, Annoni J-M, Chtioui H, Dao K, Fabritius M, et al. Long-term effects of cannabis on brain structure. Neuropsychopharmacology. 2014;39:2041–8.

22. Medina KL, Nagel BJ, Tapert SF. Abnormal cerebellar morphometry in abstinent adolescent marijuana users. Psychiatry Res. 2010;182:152–9.

23. Block RI, O’Leary DS, Ehrhardt JC, Augustinack JC, Ghoneim MM, Arndt S, et al. Effects of frequent marijuana use on brain tissue volume and composition. Neuroreport. 2000;11:491–6.

24. Cohen M, Rasser PE, Peck G, Carr VJ, Ward PB, Thompson PM, et al. Cerebellar grey-matter deficits, cannabis use and first-episode schizophrenia in adolescents and young adults. Int J Neuropsychopharmacol. 2012;15:297–307.

25. Lopez-Larson MP, Bogorodzki P, Rogowska J, McGlade E, King JB, Terry J, et al. Altered prefrontal and insular cortical thickness in adolescent marijuana users. Behav Brain Res. 2011;220:164–72.

26. Filbey FM, McQueeny T, DeWitt SJ, Mishra V. Preliminary findings demonstrating latent effects of early adolescent marijuana use onset on cortical architecture. Dev Cogn Neurosci. 2015;16:16–22.

27. Jacobus J, Squeglia LM, Meruelo AD, Castro N, Brumback T, Giedd JN, et al. Cortical thickness in adolescent marijuana and alcohol users: A three-year prospective study from adolescence to young adulthood. Dev Cogn Neurosci. 2015.

28. Button KS, Ioannidis JPA, Mokrysz C, Nosek BA, Flint J, Robinson ESJ, et al. Power failure: why small sample size undermines the reliability of neuroscience. Nat Rev Neurosci. 2013;14:365–76.

29. Gronenschild EHBM, Habets P, Jacobs HIL, Mengelers R, Rozendaal N, van Os J, et al. The effects of FreeSurfer version, workstation type, and Macintosh operating system version on anatomical volume and cortical thickness measurements. PloS One. 2012;7:e38234.

30. Mackenzie-Graham AJ, Van Horn JD, Woods RP, Crawford KL, Toga AW. Provenance in neuroimaging. NeuroImage. 2008;42:178–95.

31. Calkins ME, Merikangas KR, Moore TM, Burstein M, Behr MA, Satterthwaite TD, et al. The Philadelphia Neurodevelopmental Cohort: constructing a deep phenotyping collaborative. J Child Psychol Psychiatry. 2015.

32. Rosen AFG, Roalf DR, Ruparel K, Blake J, Seelaus K, Villa LP, et al. Quantitative assessment of structural image quality. NeuroImage. 2018;169:407–18.

33. Gennatas ED, Avants BB, Wolf DH, Satterthwaite TD, Ruparel K, Ciric R, et al. Age-related effects and sex differences in gray matter density, volume, mass, and cortical thickness from childhood to young adulthood. J Neurosci. 2017;37:5065–73.

34. Satterthwaite TD, Elliott MA, Ruparel K, Loughead J, Prabhakaran K, Calkins ME, et al. Neuroimaging of the Philadelphia neurodevelopmental cohort. NeuroImage. 2014;86:544–53.

35. Scott JC, Wolf DH, Calkins ME, Bach EC, Weidner J, Ruparel K, et al. Cognitive functioning of adolescent and young adult cannabis users in the Philadelphia Neurodevelopmental Cohort. Psychol Addict Behav. 2017;31:423–34.

36. Han C, McGue MK, Iacono WG. Lifetime tobacco, alcohol and other substance use in adolescent Minnesota twins: univariate and multivariate behavioral genetic analyses. Addiction. 1999;94:981–93.

37. Becker MP, Collins PF, Lim KO, Muetzel RL, Luciana M. Longitudinal changes in white matter microstructure after heavy cannabis use. Dev Cogn Neurosci. 2015;16:23–35.

38. Meier MH, Caspi A, Ambler A, Harrington H, Houts R, Keefe RSE, et al. Persistent cannabis users show neuropsychological decline from childhood to midlife. Proc Natl Acad Sci U S A. 2012;109:e2657–2664.

39. Shanmugan S, Wolf DH, Calkins ME, Moore TM, Ruparel K, Hopson RD, et al. Common and dissociable mechanisms of executive system dysfunction across psychiatric disorders in youth. Am J Psychiatry. 2016;173:517–26.

40. Goodkind M, Eickhoff SB, Oathes DJ, Jiang Y, Chang A, Jones-Hagata LB, et al. Identification of a common neurobiological substrate for mental illness. JAMA Psychiatry. 2015;72:305–15.

41. Tustison NJ, Avants BB, Cook PA, Zheng Y, Egan A, Yushkevich PA, et al. N4ITK: Improved N3 Bias Correction. IEEE Trans Med Imaging. 2010;29:1310–20.

42. Tustison NJ, Cook PA, Klein A, Song G, Das SR, Duda JT, et al. Large-scale evaluation of ANTs and FreeSurfer cortical thickness measurements. NeuroImage. 2014;99:166–79.

43. Avants BB, Tustison NJ, Wu J, Cook PA, Gee JC. An open source multivariate framework for n-tissue segmentation with evaluation on public data. Neuroinformatics. 2011;9:381–400.

44. Avants BB, Tustison NJ, Song G, Cook PA, Klein A, Gee JC. A Reproducible Evaluation of ANTs Similarity Metric Performance in Brain Image Registration. NeuroImage. 2011;54:2033–44.

45. Das SR, Avants BB, Grossman M, Gee JC. Registration based cortical thickness measurement. NeuroImage. 2009;45:867–79.

46. Marcus DS, Wang TH, Parker J, Csernansky JG, Morris JC, Buckner RL. Open Access Series of Imaging Studies (OASIS): cross-sectional MRI data in young, middle aged, nondemented, and demented older adults. J Cogn Neurosci. 2007;19:1498–507.

47. Wang H, Suh JW, Das SR, Pluta JB, Craige C, Yushkevich PA. Multi-Atlas Segmentation with Joint Label Fusion. IEEE Trans Pattern Anal Mach Intell. 2013;35:611–23.

48. Avants BB, Tustison NJ, Wu J, Cook PA, Gee JC. An open source multivariate framework for n-tissue segmentation with evaluation on public data. Neuroinformatics. 2011;9:381–400.

49. Gilman JM, Kuster JK, Lee S, Lee MJ, Kim BW, Makris N, et al. Cannabis use is quantitatively associated with nucleus accumbens and amygdala abnormalities in young adult recreational users. J Neurosci. 2014;34:5529–38.

50. Fischl B. FreeSurfer. NeuroImage. 2012;62:774–81.

51. Altman DG, Bland JM. How to obtain the P value from a confidence interval. BMJ. 2011;343:d2304. doi:10.1136/bmj.d2304.

52. Lakens D. Equivalence tests. Soc Psychol Personal Sci. 2017;8:355–62.

53. Wilkinson GS, Robertson GJ. WRAT 4: Wide Range Achievement Test; professional manual. Psychological Assessment Resources, Incorporated; 2006.

54. Thayer RE, YorkWilliams S, Karoly HC, Sabbineni A, Ewing SF, Bryan AD, et al. Structural neuroimaging correlates of alcohol and cannabis use in adolescents and adults. Addict Abingdon Engl. 2017;112:2144–54.

55. French L, Gray C, Leonard G, Perron M, Pike GB, Richer L, et al. Early cannabis use, polygenic risk score for schizophrenia and brain maturation in adolescence. JAMA Psychiatry. 2015;72:1002–11.

56. Cheetham A, Allen NB, Whittle S, Simmons JG, Yücel M, Lubman DI. Orbitofrontal volumes in early adolescence predict initiation of cannabis use: A 4-year longitudinal and prospective study. Biol Psychiatry. 2012;71:684–92.

57. Vijayakumar N, Allen NB, Youssef G, Dennison M, Yücel M, Simmons JG, et al. Brain development during adolescence: A mixed-longitudinal investigation of cortical thickness, surface area, and volume. Hum Brain Mapp. 2016;37:2027–38.

58. Koenders L, Cousijn J, Vingerhoets WAM, van den Brink W, Wiers RW, Meijer CJ, et al. Grey matter changes associated with heavy cannabis use: A longitudinal smri study. PLoS ONE. 2016;11.

59. Jackson NJ, Isen JD, Khoddam R, Irons D, Tuvblad C, Iacono WG, et al. Impact of adolescent marijuana use on intelligence: Results from two longitudinal twin studies. Proc Natl Acad Sci. 2016;113:E500–8.

60. Gao Y, Riklin-Raviv T, Bouix S. Shape analysis, a field in need of careful validation. Hum Brain Mapp. 2014;35:4965–78.

61. Jones DK, Knösche TR, Turner R. White matter integrity, fiber count, and other fallacies: the do’s and don’ts of diffusion MRI. NeuroImage. 2013;73:239–54.

## Supplementary References

1. Fischl B. FreeSurfer. NeuroImage. 2012;62:774–81.

2. Dale AM, Fischl B, Sereno MI. Cortical surface-based analysis. I. Segmentation and surface reconstruction. NeuroImage. 1999;9:179–94.

3. Desikan RS, Ségonne F, Fischl B, Quinn BT, Dickerson BC, Blacker D, et al. An automated labeling system for subdividing the human cerebral cortex on MRI scans into gyral based regions of interest. NeuroImage. 2006;31:968–80.

